# Human activities impact microbial communities of Amazon Mangroves

**DOI:** 10.1101/2022.08.03.501699

**Authors:** Gleyciane Machado da Costa, Sávio Souza Costa, Rafael Azevedo Barauna, Bruno Pureza Castilho, Izabel Cruz Pinheiro, Artur Silva, Ana Paula Schaan, Ândrea Kely Campos Ribeiro-dos-Santos, Diego Assis das Graças

**Affiliations:** Laboratory of Genomics and Bioinformatics, Center of Genomics and Systems Biology, Institute of Biological Sciences, Federal University of Pará, Belém, Brazil; Laboratory of Biological Engineering, Guamá Science and Technology Park, Belém, Brazil; Federal University of Pará, Belém, Brazil; Laboratory of Medical and Human Genetics, Institute of Biological Sciences, Federal University of Pará, Belém, Brazil

**Keywords:** Amazon, Microbiota Mangroves, Metagenomics, Anthropogenic Action, Microbial Diversity

## Abstract

Mangroves provide a unique ecological environment for complex microbial communities. They are particularly important in controlling the chemical environment of the ecosystem. Microbial diversity analyses of these ecosystems help us understand the environmental dynamics and the changes due to impacts. Amazonian mangroves are distributed across Amapá, Pará and Maranhão states occupying an area of 9,000 km^2^ and correspond to 70% of the mangroves in Brazil, in which studies on microbial biodiversity are extremely rare. The present study aimed to determine changes in microbial community structure along the PA-458 highway that divided the mangrove. Mangrove samples were collected from three zones, (i) under anthropogenic actions, (ii) in the process of recovery and (ii) preserved. Total DNA was extracted, submitted for 16S rDNA amplification and sequenced on the Illumina MiSeq platform. Reads were analyzed using USEARCH tools and packages available in R Platform. The most abundant phyla were Proteobacteria, followed by Firmicutes and Bacteroidetes in all three mangrove locations. Genera such as *Desulfuromonas*, *Desulfatiglans*, *Collinsella*, *Dorea*, *Dialister* were not found in the impacted mangrove. The impacted mangrove proved to be ecologically different from other mangroves. Our results show that human impact in the mangrove areas has resulted in loss of biodiversity, caused by the construction of the PA-458 highway, reducing the support and maintenance of habitat life and the ecosystem’s resilience to disturbance.

## • Introduction

Mangroves are one of the most dynamic and productive ecosystems and have great ecological importance. Located along tropical and subtropical regions, they constitute more than half of the terrestrial coastline (Sahoo, Dhal, 2009). In the Amazon, they occupy an area of 9,000 km^2^, but studies on the microbial biodiversity of these ecosystems are extremely scarce. These mangroves have trophic networks of great complexity, which makes them ideal habitats to harbor great microbial diversity and thus maintain essential functions for the maintenance of the ecosystem (Gomes et al., 2010; dos Santos et al., 2011).

Microbial diversity is strongly influenced by biogeographic, anthropogenic and ecological properties, including nutrient availability and the presence of organic and inorganic compounds (Ghizelini et al., 2012). Salinity is a particularity of this biome and is related to the proximity to the sea and the variation of the tide, which promotes frequent anaerobic conditions (Holguin et al., 2001; Ferreira et al., 2010).

The degradation of organic matter and pollutants, in addition to recycling and providing nutrients to other organisms, are microbial processes of great importance in mangroves (Dias et al., 2012). Nitrogen and phosphorus are often limiting nutrients for microbial growth in various environments. Variations in phosphorus concentrations in mangroves, associated with anthropogenic impacts, can lead to high susceptibility to eutrophication events. However, the availability of inorganic and organic phosphorus, as well as their microbial cycling dynamics in this ecosystem are still poorly understood.

The Bragança Region is located on the Atlantic coast, approximately 200 km east of the nearest capital Belém, in the state of Pará, Brazil. Its mangrove peninsula encompasses the estuaries of the Caeté and Tapera-Açu rivers and is situated in the second largest continuous mangrove ecosystem in the world (Kjerfve et al., 1997).

The construction of the PA-458 highway caused the destruction of mangrove areas in the Ajuruteua Peninsula (Lara, Cohen, 2002), which affects not only aquatic systems that depend on the exchange of nutrients (Fernandes et al., 2015), but also animal nutrition and the riverine subsistence economy (Fernandes, 2003). Studies of the microbial communities in mangroves become especially important since little is known about the biodiversity of these environments, even more so in the Amazon. In this study, we describe and compare changes in the microbial community structures in impacted and non-impacted mangroves by the construction of the PA-458 highway in the municipality of Bragança in northeastern Pará, using an amplicon sequencing approach.

- Materials and Methods
- Sample Collection

Mangrove samples were collected in triplicate at six points along the PA-458 highway that connects the municipality of Bragança to the coastal village of Ajuruteua (Fig 1). The highway is in the northeast region of the State of Pará, Brazil, and collection points were separated in East and West side of the highway, totaling 18 samples.

**Figura 1.**
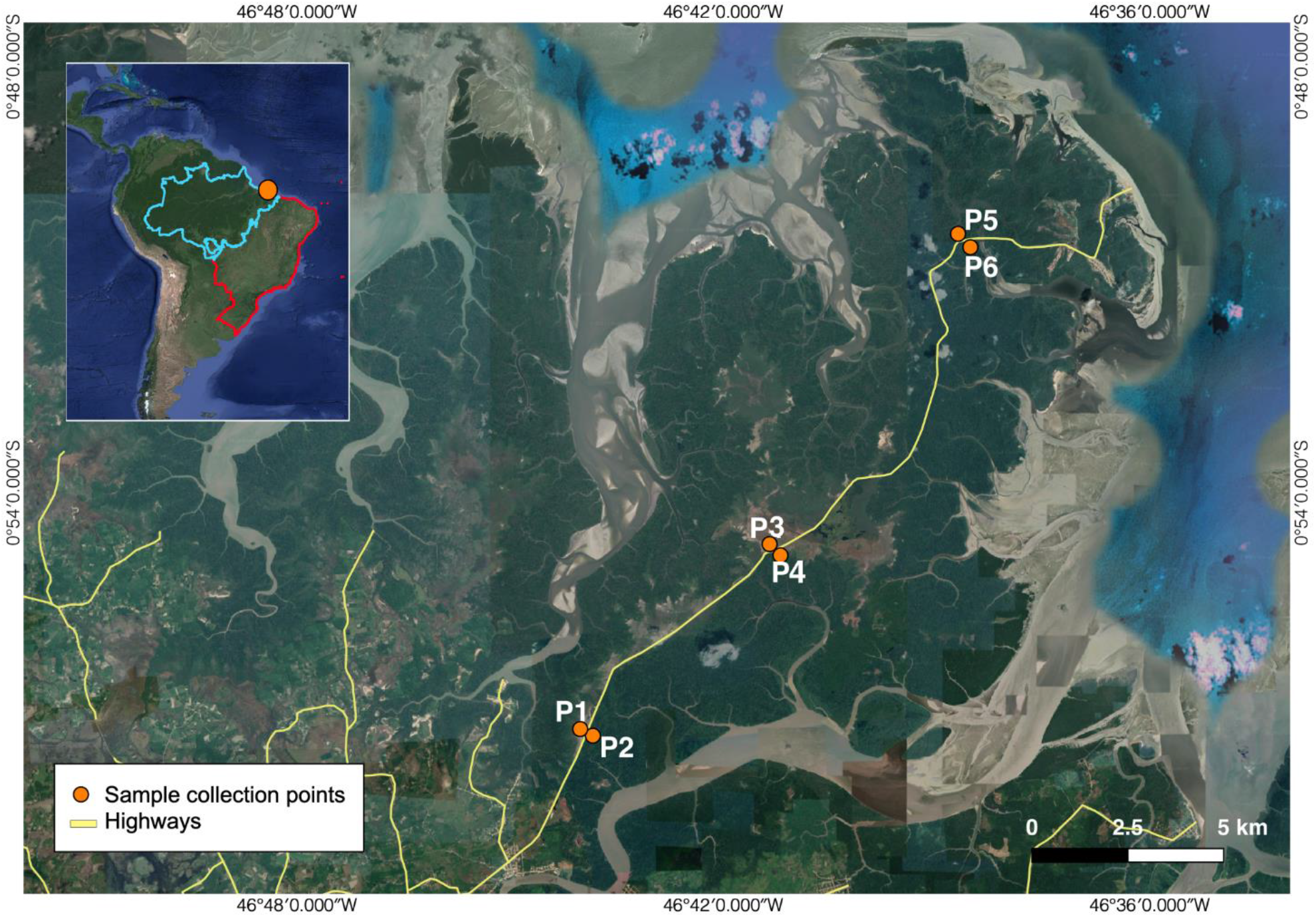
Location of each sampling site along highway PA-458 in relation to Brazil (red line) and the Brazilian Amazon territory (blue line).

All three collection points on the East side of the highway were considered preserved and labeled as P2, P4 and P6. On the West side, samples P1, P3 and P5 were respectively classified as under the effect of anthropogenic impacts, in the process of recovery and preserved. At each site, approximately 30 g of subsurface sediment (~30 cm depth) was collected and immediately transferred to a 50 ml tube. Samples were chilled on ice until arrival at the laboratory and then stored at −80°C until further procedures.

### • DNA Extraction and Sequencing of 16S rRNA genes

Total DNA from mangrove samples was extracted using the Powersoil DNA Isolation Kit (MO BIO, Carlsbad, CA USA) according to the manufacturer’s protocol with small modifications. Eluted DNA was quantified with fluorometry and subsequently stored at −20 °C. Library preparation was carried out according to the Illumina Metagenomic Sequencing Library Prep protocol with established primers and Illumina Nextera adapters targeting the V3-V4 region of the 16S rDNA, as follows: Bakt_341F (CCTACGGGNGGCWGCAG) e Bakt_805R (GACTACHVGGGTATCTAATCC). Libraries were subsequently pooled and quantified with TapeStation (Agilent, Santa Clara, CA) prior to sequencing on the Illumina MiSeq platform (Figure 2).

**Figure 2.**
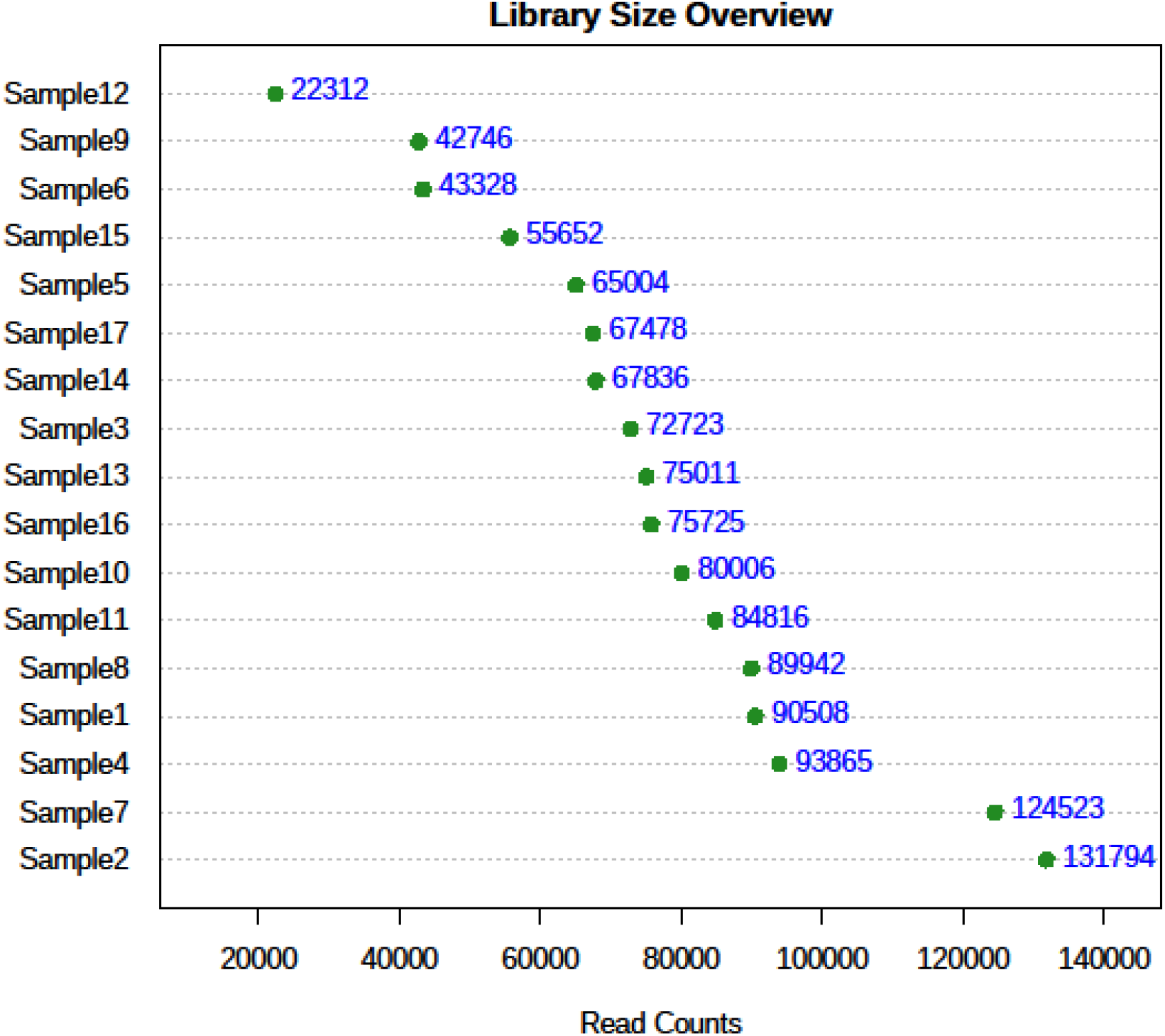
Overview of the number of reads for each sample after sequencing.

### • Data Analysis

After sequencing, low-quality reads (maximum probability of error 0.5), smaller than 200 bp, and chimeric artifacts were removed using USEARCH (Edgar, 2010). Sequences were clustered into Operational Taxonomic Units (OTU) at 97% similarity using USEARCH. Taxonomy was assigned to each OTU performing BLAST searches in the SILVA v.132 database with a maximum E value of 1e-5 (Quast, 2013).

Alpha diversity, rarefaction curves and relative abundance indices of bacterial classification at the phyla and genus level among the sample groups were calculated and determined using the phyloseq (Mcmurdie, Holmes, 2013), vegan (Oksanen, 2017) and ggplot2 (Wickham, 2016) packages implemented in R (R project). Furthermore, beta diversity was visualized using a non-metric multidimensional scaling (NMDS) analysis based on Bray-Curtis dissimilarity indexes extracted from genus level taxonomies using the vegan package, implemented in R.

### • Physico-chemical Parameters

The analysis of the following physico-chemical parameters: Carbon (C), Organic Matter (OM), Total Nitrogen (N), Phosphorus (P), Potassium (K), Sodium (Na), Aluminum (Al), Calcium (Ca), Magnesium (Mg), pH and Potential Acidity (H+Al) were carried out at the Brazilian Agricultural Research Corporation (EMBRAPA).

## Results

The data presented in [Table 1] correspond to the results of the physico-chemical analyses of mangrove sediment samples. We observed a difference in the amount of carbon, organic matter, phosphorus, potassium and cation-exchange capacity (CEC) when comparing the mangrove sediment impacted by anthropogenic actions and preserved (P1 and P2).

**Table 1.** Physicochemical parameters table.

| Samples | C | OM | N | N | Relaç | P | K | Na | Al | Ca | Ca+Mg | pH | H+Al | CTC |  | Saturation |  |
| --- | --- | --- | --- | --- | --- | --- | --- | --- | --- | --- | --- | --- | --- | --- | --- | --- | --- |
| ID | g/kg | g/kg | % | g/kg | C/N | mg/dm<sup>3</sup> |  |  | cmol/dm<sup>3</sup> |  |  | Water | cmolc/dm<sup>3</sup> | Total (cmol/dm<sup>3</sup>) | Effective (cmol/dm<sup>3</sup>) | Base (V%) | Aluminum (m%) |
| P1 | 10,8 | 18,7 | 0,08 | 0,81 | 13,3 | 91 | 1303 | 23072 | 0 | 5 | 33,2 | 5,9 | 0 | 136,81 | 136,83 | 100 | 0,01 |
| P2 | 16,5 | 28,5 | 0,07 | 0,73 | 22,8 | 16 | 444 | 3957 | 0,1 | 2,5 | 8,5 | 6,4 | 3,82 | 30,66 | 26,91 | 87,55 | 0,26 |
| P3 | 10,4 | 18 | 0,1 | 1,03 | 10,1 | 85 | 1391 | 19967 | 0 | 6,1 | 29,9 | 5,6 | 0,49 | 120,75 | 120,29 | 99,6 | 0,02 |
| P4 | 24,8 | 42,8 | 0,13 | 1,29 | 19,2 | 43 | 788 | 10957 | 0 | 3,6 | 17,4 | 5,8 | 2,09 | 69,14 | 67,07 | 96,97 | 0,04 |
| P5 | 21,7 | 37,5 | 0,07 | 0,73 | 30 | 34 | 919 | 11976 | 0 | 3,2 | 15,7 | 5,1 | 0,79 | 70,93 | 70,18 | 98,89 | 0,04 |
| P6 | 8,4 | 14,5 | 0,07 | 0,7 | 11,9 | 28 | 876 | 12082 | 0 | 3,1 | 15,2 | 5,5 | 0,9 | 70,89 | 70,03 | 98,73 | 0,04 |

The rarefaction curve reveals that the samples from the environment with anthropogenic impact showed lower species richness compared to the others, obtaining an average of 2,000 OTUs, while preserved samples showed 2,500-6,000 OTUs, and samples from the recovery area 4,000-7,000 OTUs (Fig 3).

**Figure 3.**
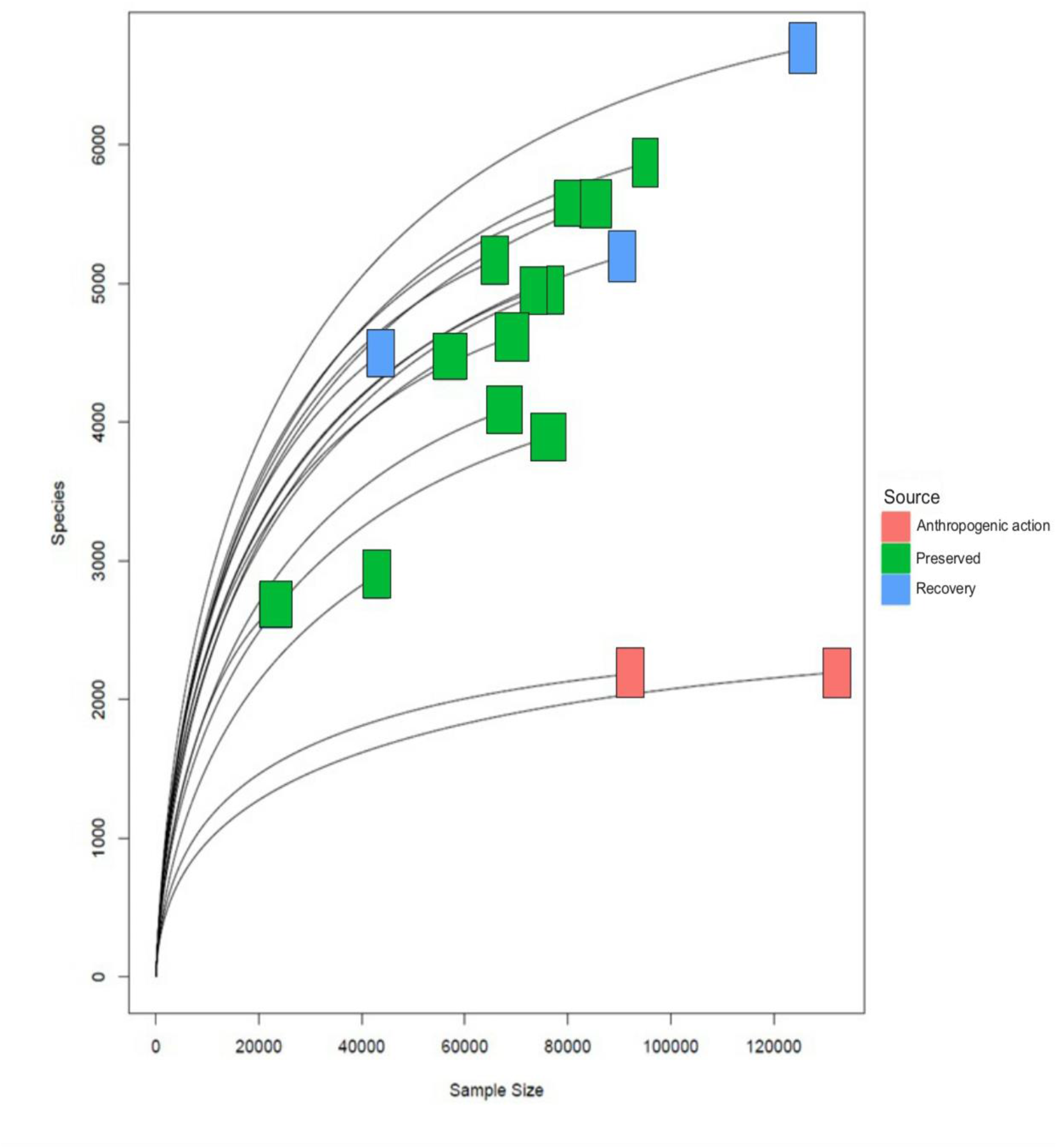
Rarefaction curve of the samples in the three mangrove environments, showing the number of sequenced reads (x axis) by the number of observed species.

Alpha diversity results of the samples collected in the three environments reveal the existence of a discrepant difference in microbial diversity between the environment impacted by anthropogenic actions and both the preserved and the recovery samples, for which the former has lower alpha diversity metrics as observed with the Chao1, Shannon, and Simpson diversity index (Fig 4). These results confirm that anthropogenic activities in the mangrove have reduced microbial diversity in the environment.

**Figure 4.**
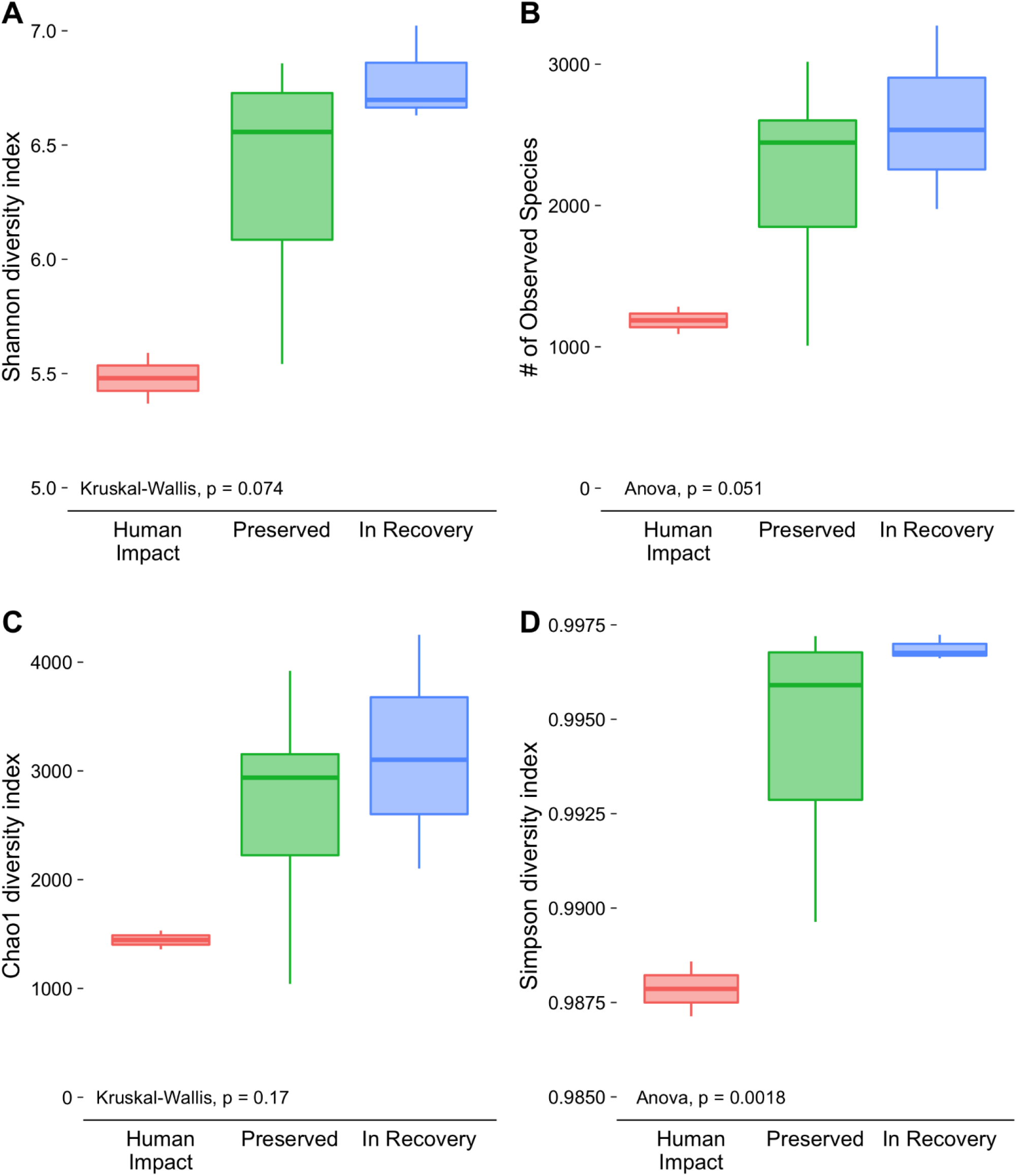
Chao1, Shannon, and Simpson diversity index of mangroves under anthropogenic action, in the process of recovery and preserved.

The NMDS analysis showed that samples under anthropogenic action are different from the preserved and the recovery sediment (Fig 5). This finding is also supported by the taxonomic composition analysis of the samples.

**Figure 5.**
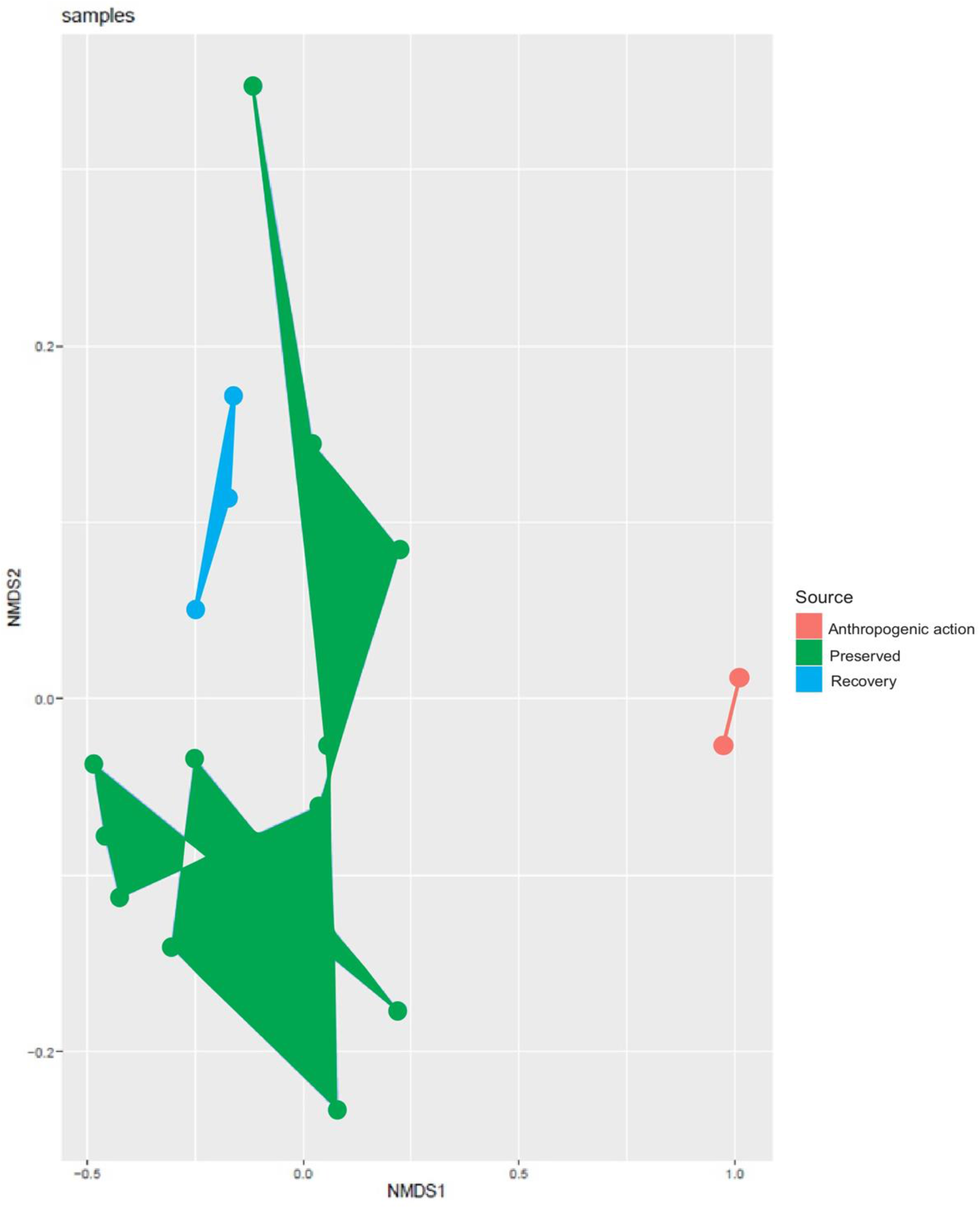
Non-Metric Multidimensional Scaling (NMDS) showing the ecological geometric distance between mangrove environments.

The most abundant phyla were Proteobacteria, followed by Firmicutes and Bacteroidetes in all three mangrove locations. The Bacteroidetes phylum was observed in higher abundance in the location impacted by human activity when compared to collection points (Fig. 5). Further, in this location we observe lower abundance of Acidobacteria and a smaller amount of different genera (Fig. 7). Beta diversity, assessed through Bray-Curtis distances, showed that samples collected in the human impacted environment have a distinct composition when compared to the preserved and in-recovery áreas, which overlapped in the Principal Coordinate Analysis (Fig. 8).

**Figure 6.**
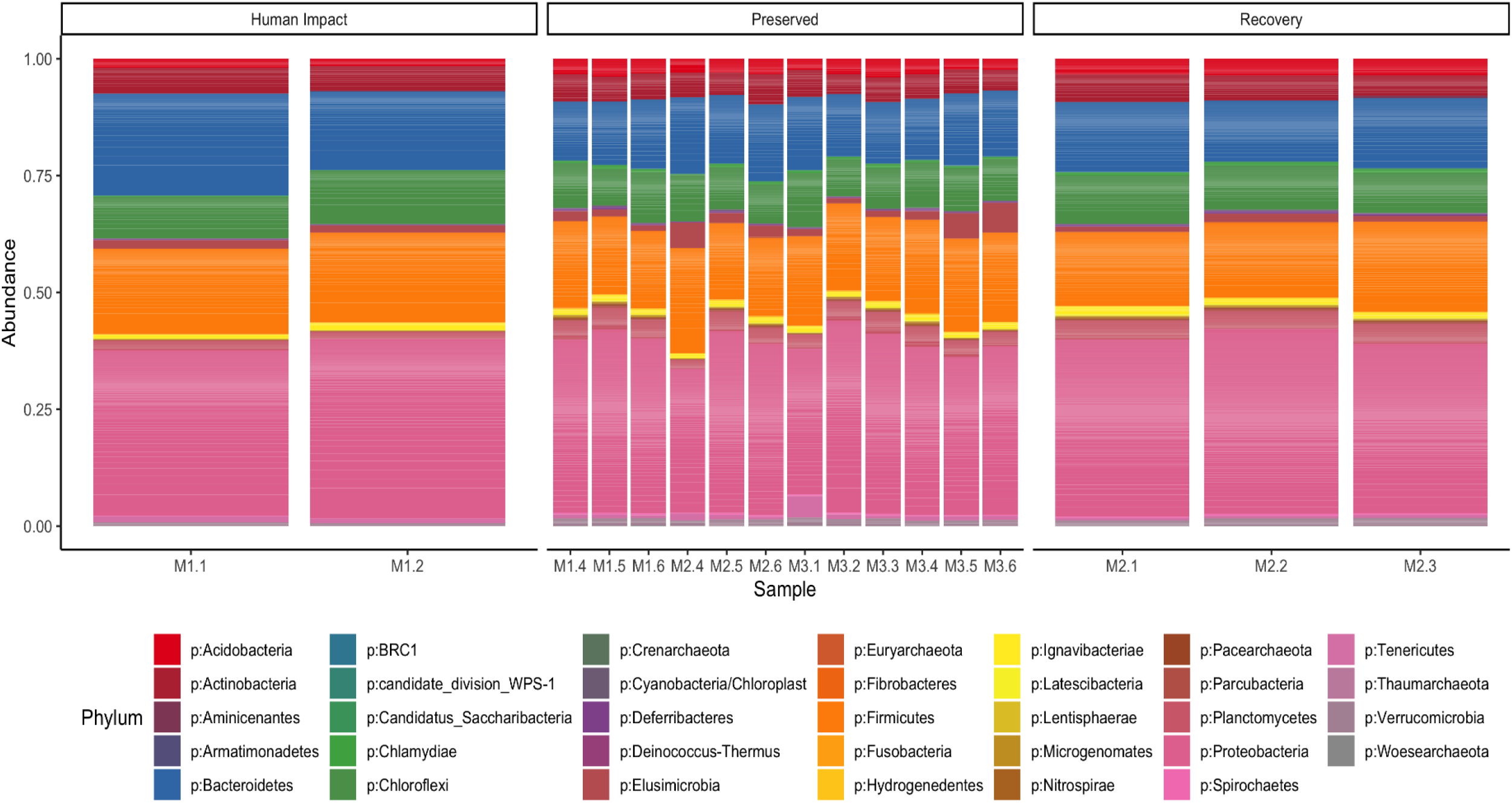
Relative abundance graph at phylum level in mangrove environments.

**Figure 7.**
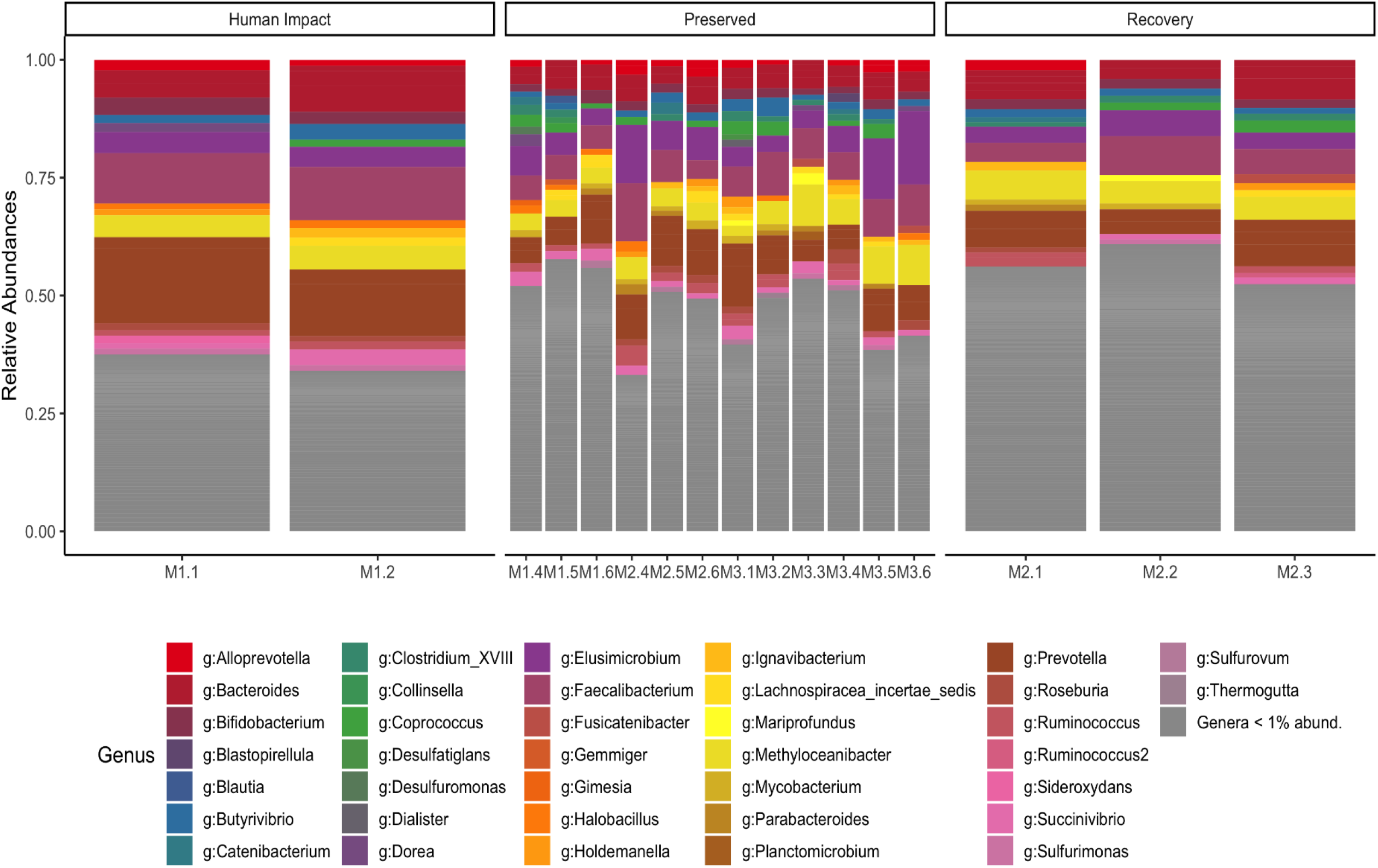
Relative abundance graph at genus level in mangrove environments.

**Figure 8.**
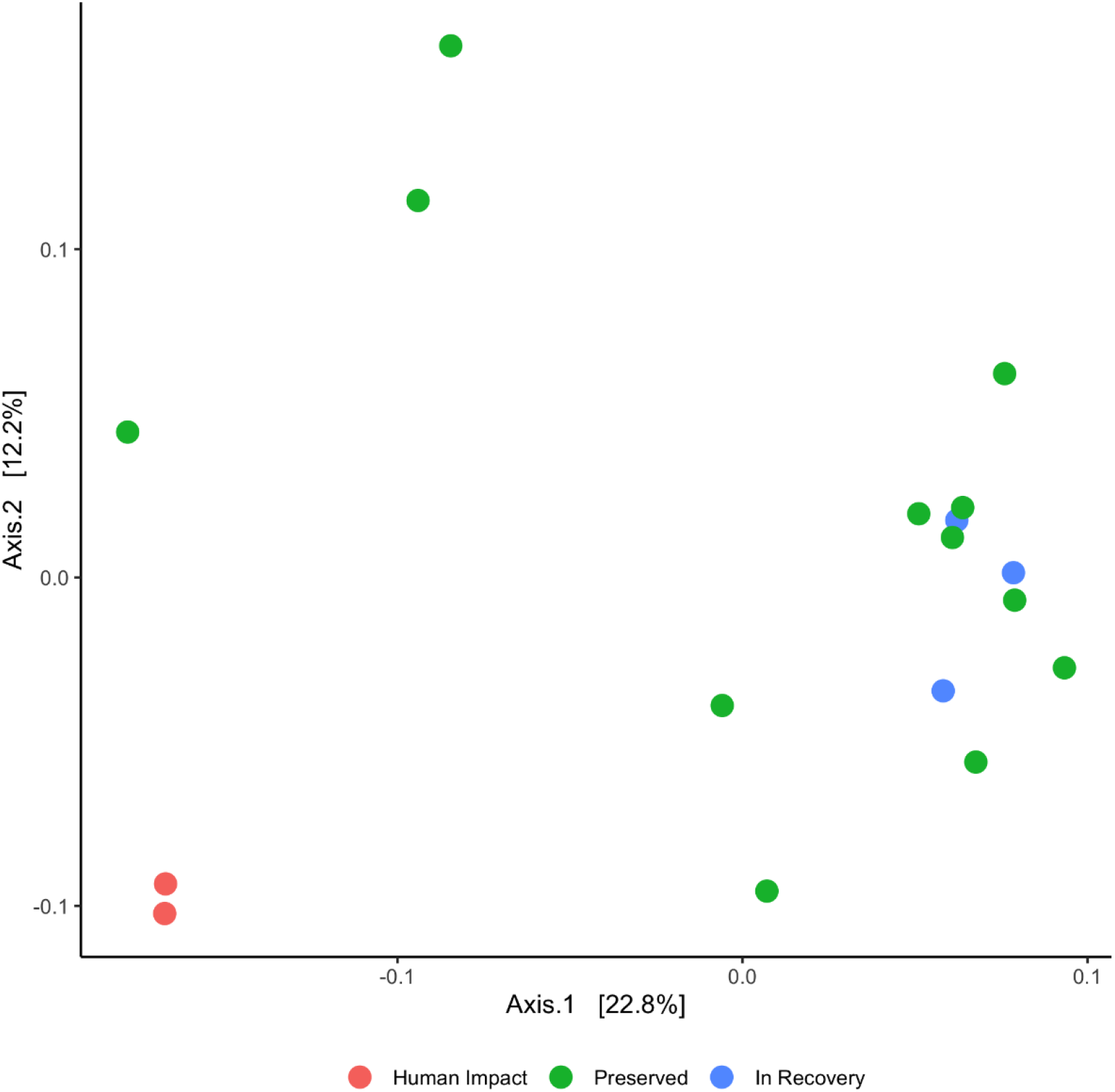
Principal Coordinate Analysis (PCoA) of Bray-Curtis distances between different collection points in the Amazonian Mangrove.

## 1 Discussion

Among the species most found in Brazilian mangroves, 53-60% belong to the phylum Proteobacteria (Nogueira et al., 2015), which correspond to about half of the bacterial genomes recorded at the National Center for Biotechnology Information (NCBI), corroborating the results of this study (Taketani et al., 2018; Haldar, Nazareth, 2018; Wu et al., 2016; Fernandes et al., 2015; Gong et al., 2019). Represented mostly by the classes of Deltaproteobacteria and Gammaproteobacteria, Proteobacteria are part of the core microbiome of mangroves worldwide (Cotta et al., 2019; Luis et al., 2019; Sadaiappan et al., 2018). Proteobacteria are found essentially in freshwater and marine environments in all latitudes, where they are responsible for the degradation of organic matter (Polyakov et al., 2009).

In a study carried out in three mangrove areas in the state of São Paulo (contaminated by oil, under anthropogenic effect and preserved), more than 45% of all microbial diversity was represented by Proteobacteria in all three areas. On the other hand, the same study showed a percentage below 15% for Firmicutes, and even lower abundances for Bacteroidetes in mangrove areas under the effect of anthropogenic action, despite the former being reported as the second most abundant phylum after Proteobacteria, and followed by Actinobacteria and Bacteroidetes (Andreote et al., 2012). In our observations, the Firmicutes phylum showed the same proportions in all mangrove environments, which may be due to the small sample size.

A lower predominance of the phyla Bacteroidetes was also found in contaminated mangroves on the island of Hainan in China, but the phylum Actinobacteria was the most abundant after the phylum Proteobacterias (Li et al., 2019). In contaminated mangroves along the Red Sea Coast in Saudi Arabia, Bacteroidetes also showed a lower predominance, however being the second most abundant phylum behind Proteobacterias (Ullah et al., 2017).

In all three studies discussed here (Andreote et al., 2012, Li et al., 2019 and Ullah et al., 2017), when the phyla of the impacted mangrove areas are compared with pristine zones, there is an increase in the Firmicutes phylum in the impacted mangrove areas, which did not occur in our study, the Firmicutes and Chloroflexi phylum remained basically in the same proportion. It is worth mentioning that the mangrove areas impacted in the studies discussed here had a certain amount of vegetation in their environments, which was not the case for our mangrove study area under anthropogenic action. A study of the vertical distributions of rhizosphere bacteria from three mangrove species in Beilun Estuary located in southern China revealed that plant species and depth played a role in the formation of the microbial communities (Wu et al., 2016), factors that were not considered here.

The underexplored phylum Acidobacteria represent a group of soil bacteria, whose members are widely and evenly distributed throughout almost all ecosystems (Mushinski et al., 2018). These bacteria may have an important contribution in the mangrove ecosystem, as they are phylogenetic diversity and may play many roles in various biogeochemical cycles while being metabolic versatility (Naether et al., 2012). In the present study, the Acidobacteria phylum showed the greatest difference between the three mangrove environments. We observed lower proportions in the anthropogenically impacted mangrove, which is expected for an area with biodiversity loss (Kielak et al., 2016a, Crits-Christoph et al., 2018)

At the genus level, we observed differential distribution patterns in the impacted and non-impacted mangrove regions, including the loss of some genera in the impacted areas. Here, we consider the absence of these bacterial groups in the impacted environment as indicative as the result of human interference. Such bacterial groups may represent transient genera that play important ecological processes of maintenance in the mangrove ecosystem (e.g. *Desulfuromonas, Desulfatiglans, Collinsella, Dorea, Dialister*).

The Desulfuromonas and Desulftiglans genera are sulfate-reducing bacteria that are involved in the sulfur cycling process, one of the main processes that occur within the mangrove ecosystem. As an anaerobic environment, rich in sulfate and organic matter, sulfate-reducing bacteria play an important role in helping to maintain the high productivity of this ecosystem, acting as primary decomposers of organic matter (Alongi et al., 1993). There was also an increase in Bacteroidetes which corroborates with human impacts on the mangrove.

The significance of this study is to show that human factors have an influence on microbial diversity and possibly the function of mangrove ecosystems. We showed that in mangrove sediments under anthropogenic action there was a loss of bacterial diversity when compared to preserved and recovering mangroves, which may indicate that contamination in mangrove sediments may have a negative effect on nutrient decomposition, hindering overall microbial communities in the habitat.

## 2 Conflict of Interest

The authors declare that the research was conducted in the absence of any commercial or financial relationships that could be construed as a potential conflict of interest.

### • Author Contributions

GC and DG: analyzed, interpreted the data, and wrote the manuscript. SC, RB, AS: contributed to the bioinformatics analyses. BC and IP: collected the samples. APS and ÂR-d-S: contributed to the sequencing analysis. DG: conceived the study.

### • Funding

This work was supported by the Coordenação de Aperfeiçoamento de Pessoal de Nível Superior (CAPES) and Fundação Amazônia de Amparo a Estudos e Pesquisas (FAPESPA)

